# Three-level Sleep Stage Classification Based on Wrist-worn Accelerometry Data Alone

**DOI:** 10.1101/2021.08.10.455812

**Authors:** Jian Hu, Haochang Shou

## Abstract

**Objective:** The use of wearable sensor devices on daily basis to track real-time movements during wake and sleep has provided opportunities for automatic sleep quantification using such data. Existing algorithms for classifying sleep stages often require large training data and multiple input signals including heart rate and respiratory data. We aimed to examine the capability of classifying sleep stages using sensible features directly from accelerometers only with the aid of advanced recurrent neural networks.

**Materials and Methods:** We analyzed a publicly available dataset with accelerometry data in 5s epoch length and polysomnography assessments. We developed long short-term memory (LSTM) models that take the 3-axis accelerations, angles, and temperatures from concurrent and historic observation windows to predict wake, REM and non-REM sleep. Leave-one-subject-out experiments were conducted to compare and evaluate the model performance with conventional nonsequential machine learning models using metrics such as multiclass training and testing accuracy, weighted precision, F1 score and area-under-the-curve (AUC).

**Results:** Our sequential analysis framework outperforms traditional non-sequential models in all aspects of model evaluation metrics. We achieved an average of 65% and a maximum of 81% validation accuracy for classifying three sleep labels even with a relatively small training sample of clinical visitors. The presence of two additional derived variables, local variability and range, have shown to strongly improve the model performance.

**Discussion:** Results indicate that it is crucial to account for deep temporal dependency and assess local variability of the features. The post-hoc analysis of individual model performances on subjects’ demographic characteristics also suggest the need of including pathological samples in the training data in order to develop robust machine learning models that are capable of capturing normal and anomaly sleep patterns in the population.

## INTRODUCTION

Disturbances in sleep and anomaly movement are known to be closely related to various clinical endpoints ^1–5^. Traditionally, sleep detection and evaluations were mostly assessed via self-reported sleep diary or using laboratory polysomnography (PSG) in sleep clinics. Sleep diary, as the most commonly used approach in recording sleep-related information such as time of bed, time of rising, and number of awakenings during sleep, relies entirely upon subjects’ recall. Hence the data are subject to recall bias and could suffer from missing information. PSG, on the other hand, are often used as the gold standard for objectively diagnosing sleep disorders. PSG uses a combination of electroencephalographic (EEG), electrooculographic, and electromyographic recordings to measure detailed data of heart rate, blood oxygen, breathing and eye/leg movements, and relies on manual expert scoring to classify sleep stages ^6,7^. However, since PSGs are mostly limited within the sleep clinics and are often costly, data are rarely available for more than one day for large study populations. Meanwhile, wearable sensor devices such as accelerometers have become more and more accessible and are commonly used as alternative tools to track sleep as they are reported to be highly correlated with PSG in monitoring sleep and wakefulness^8^. Many large-scale population studies such as the UK Biobank and National Health and Nutrition Examination Study (NHANES) ^9–11^ are collecting multiple days of 24-hr movement and sleep data using accelerometers. Hence in order to objectively evaluate sleep among the general population in their free-living conditions, it has become of great interest among researchers to develop automatic algorithms that can reliably extract important features of sleep and circadian rhyme based on accelerometry data.

Data obtained from accelerometers are often continuous time series of accelerations over the three spatial dimensions (x, y, and z) with a frequency as high as 30-100Hz. They are then processed into an epoch grid of every 5-second to 1-minute resolution. Additional data such as light exposure and ambient temperature are often collected by many types of devices as well. Early methods mainly focused on detecting the binary sleep and awake cycle and often relied upon arbitrary cut-points using one or a few accelerometry features^12^. Van Hees et al. ^13^ proposed a novel method for detecting sleep period time (SPT) that was based on thresholding the change of the local variabilities of estimated arm angles (z-angle). Several recent works explored machine learning (ML) methods such as hidden Markov models (HMM) and neural networks for classifying sleep and awake from continuous actigraphy measures ^14,15^. In particular, recurrent neural networks (RNN) are a class of sequential neural networks that could process a sequence of input features with retained temporal order. Thus, they are able to remember the historical information while predicting the current state. Taking advantage of time series information in such an efficient way, sequential neural networks are expected to achieve better performance compared to the non-sequential models. In addition to the standard RNN, the Long Short Term Memory Network (LSTM) is a special variant of RNN which utilizes a 'memory cell' that could maintain information in memory for long periods of time. Compared to standard RNN, LSTM is able to learn longer-term dependencies and is proved to show superior performance in classifying dependent states in many studies. Several recent works have shown success in predicting sleep status using ‘deep temporal modeling’ such as LSTM by taking a comprehensive set of features from PSG or heart rate data from electrocardiography (ECG) ^16–18^.

In this paper, we explore the ability of actigraphy in classifying beyond the binary sleep/awake status by utilizing a full set of accelerometry features and the most advanced machine learning models over relatively limited sample size. In particular, we aim to differentiate both wake and sleep, and further rapid eye movement (REM) and non-REM during sleep using features such as 3-axis accelerations, wrist angles and local temporal variations. **Figure 1** summarizes the general workflow of feature extraction and constructing an LSTM network for our data. We constructed several sequential neural networks that are tailored for time series data and compared their performance with other competing models. Although there exist a few recent studies that classified sleep stages using sensor data achieving close to 80% accuracies^14,16,17^, most of them required inputs from multiple devices measuring various biological signals including respiratory rate, heart rate, and limb movements. Moreover, the models typically need to be trained on large cohort samples with several thousand participants. Hence such algorithms might not be directly applicable to many clinical studies where only data from a single wearable device such as accelerometers are collected at home. Unlike the other published papers, the goal of our paper is not to produce a classification algorithm that outperforms all the existing ones. Instead, we would like to assess whether machine learning models can achieve satisfactory performance in differentiating sleep stages with features extracted solely from wrist-worn accelerometers and when only limited training data are available.

**Figure 1.**
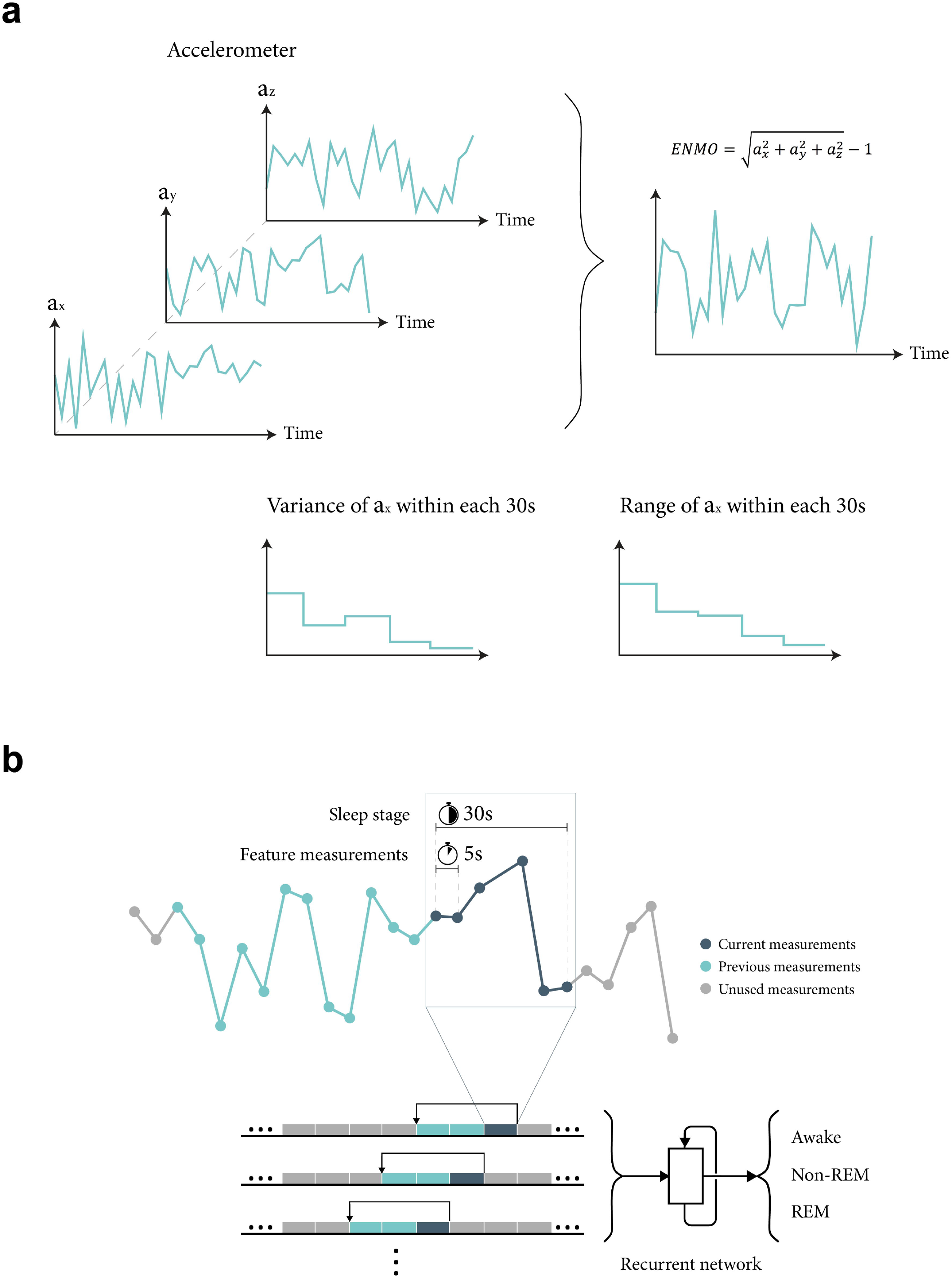
Workflow. **a), Feature extraction.** ENMO is calculated using the median values of raw acceleration in x, y and z coordinates within each 5-second epoch; variance and range of the measurements within each 30 seconds interval are calculated as artificial features. **b), Create sequential input with prior intervals.** Measurements from the wrist-worn devices are recorded every 5 seconds and the sleep stage is evaluated every 30 seconds. The past features from a fixed number of prior intervals are fed into the recurrent model subsequently.

## RESULTS

### 1. Binary Classification: Awake and Asleep

As a proof of concept, we first conduct binary classification to differentiate time at sleep versus awake. In total, our data contains 36,324 sleep state records from 28 participants, among which 12,360 are asleep and the remaining 23,964 are awake. A naïve model which gives all prediction as awake will achieve an accuracy of 69.46% and is treated as the reference baseline for comparison. We compare RNN and LSTM models with several ML classifiers such as logistic regression, perceptron, extra tree, and random forest that have previously been reported^19,20^ to achieve good performance in similar tasks. Note that since neither of the aforementioned conventional ML models was originally designed for time series data as in our data, we treat the sequential measures within each 30-second window as independent features. For a fair comparison, the RNN and LSTM were also trained using concurrent information as input without considering the historical information. The model performances are assessed via training and validation accuracy, precision, recall and weighted F1 score in the test data. Area-under-curve (AUC) was also calculated for logistic regression, RNN and LSTM that produce continuous probability estimation.

The performance of all the models is summarized in **Table 1**. Given that the outcomes are observed to be highly imbalanced, models such as Extra Tree may overfit the data and tend to predict all the records as awake. Among all of the non-sequential models, logistic regression achieves the highest validation accuracy of 73.1%. Since our dataset only contains variables from patient characteristics and measurements from the wrist-worn devices, these four models’ performance was inferior to those reported in Palotti et al. (2019)^19^. RNN and LSTM without previous sequential information (b=0) do not show superior performance compared to non-sequential models.

**Table 1:**
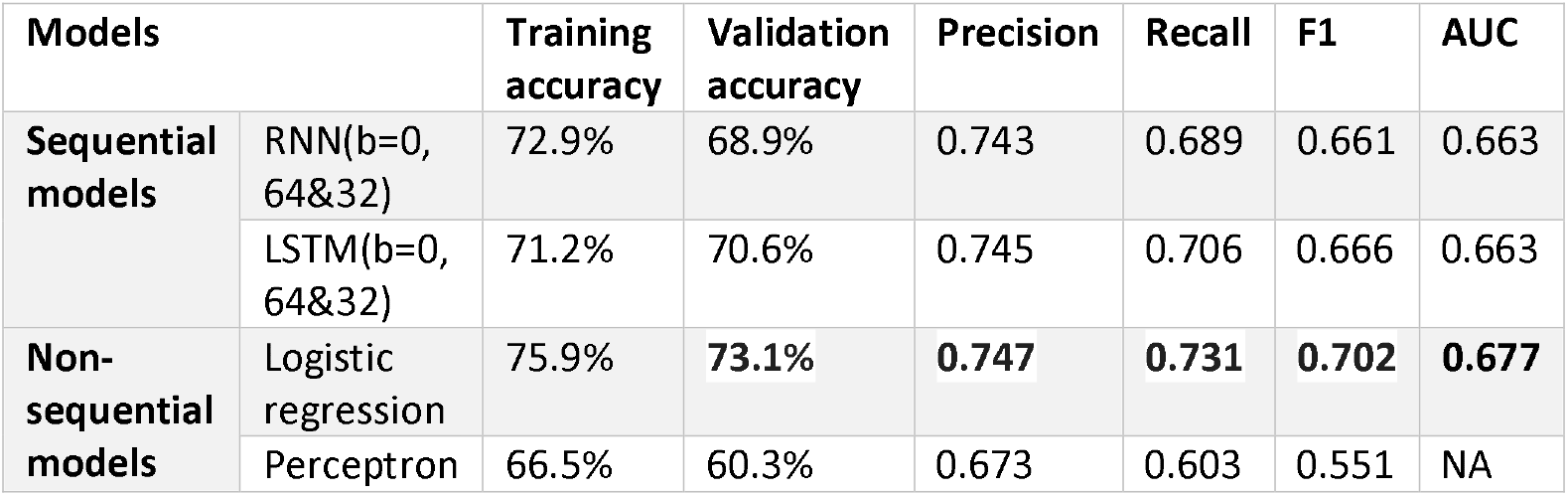
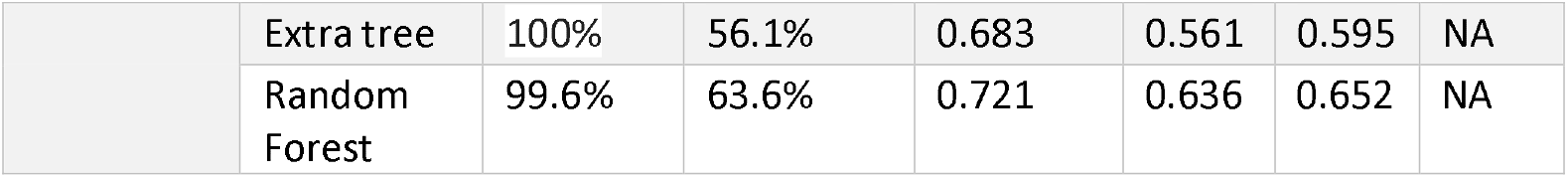
Model performance of classifying sleep vs. awake via leave-one-out experiments. The metrics were averaged across 27 rounds. RNN and LSTM were tuned to contain two hidden layers with 64 and 32 nodes each. Features were treated as independent, and no historical data was included as inputs (b=0).

| Models |  | Training accuracy | Validation accuracy | Precision | Recall | F1 | AUC |
| --- | --- | --- | --- | --- | --- | --- | --- |
| Sequential models | RNN(b=0, 64&32) | 72.9% | 68.9% | 0.743 | 0.689 | 0.661 | 0.663 |
|  | LSTM(b=0, 64&32) | 71.2% | 70.6% | 0.745 | 0.706 | 0.666 | 0.663 |
| Non-sequential models | Logistic regression | 75.9% | **73.1%** | **0.747** | **0.731** | **0.702** | **0.677** |
|  | Perceptron | 66.5% | 60.3% | 0.673 | 0.603 | 0.551 | NA |
|  | Extra tree | 100% | 56.1% | 0.683 | 0.561 | 0.595 | NA |
|  | Random Forest | 99.6% | 63.6% | 0.721 | 0.636 | 0.652 | NA |

### 2. Three-level Classification: Awake, REM and Non-REM

Our next aim is to classify the sleep stages into three levels: awake, REM and non-REM. Here the naïve model, which predicts all as non-REM, will achieve a performance of 57.7% and will be referred to as the baseline. Among the four non-sequential machine learning models evaluated in the previous section, logistic regression achieves the best overall performance with 63% validation accuracy, 0.57 F1 score and 0.66 AUC (**Supplement Table A.1**). To further improve the model performance, we investigate approaches that allow the inclusion of information from historical data in the time series. We start from an RNN with two hidden layers of 64 nodes and 32 nodes. To take the previous time-series information into account, we feed the past features from b prior intervals (each interval lasts 5 seconds) to the RNN subsequently. By doing this, RNN learns and uses sequential dependency from previous data in predicting the current sleep stage. We varied the hyper-parameter b from 0 to 40 and recorded the change of RNN’s performance (**Supplement Table A.1**). As the longer period of historical data was accounted, for all of the 6 metrics improved, with F1 score increased from 0.524 to 0.549, and AUC increased from 0.648 to 0.658. Despite that we have the advantage of RNN for sequential data, it does not show significant improvement compared to the simple logistic regressions. The reason, suggested by the low training accuracy, might be that the current model structure was too small to handle a large amount of historical information. Therefore, we enlarged the hidden layer size of RNN and assessed its performance using a similar strategy. As shown in **Supplement Table A.1**, the RNN with 2 layers of 128 and 64 nodes achieved better performance in all 6 metrics comparing to the smaller RNN and logistic regression. Also, as shown in **Figure 2a**, as parameter b increased from 0 to 80, accuracy, F1 score and AUC continued to increase. Peak performance was achieved at b=40, then plateaued and fluctuated with larger historic window sizes. In order to utilize more sequential features, we also implemented LSTM, which is suitable for incorporating longer sequential information than the standard RNN. As shown in **Figure 2b**, the average validation accuracy and F1 score of LSTM model reach the maximum value (65% accuracy and 0.609 F1) at b=50, or when prior 250 seconds of records were accounted. AUC was the highest to be 0.685 when b =80 or with 400 seconds of historical data. (**Supplement Table A.1**)

**Figure 2.**
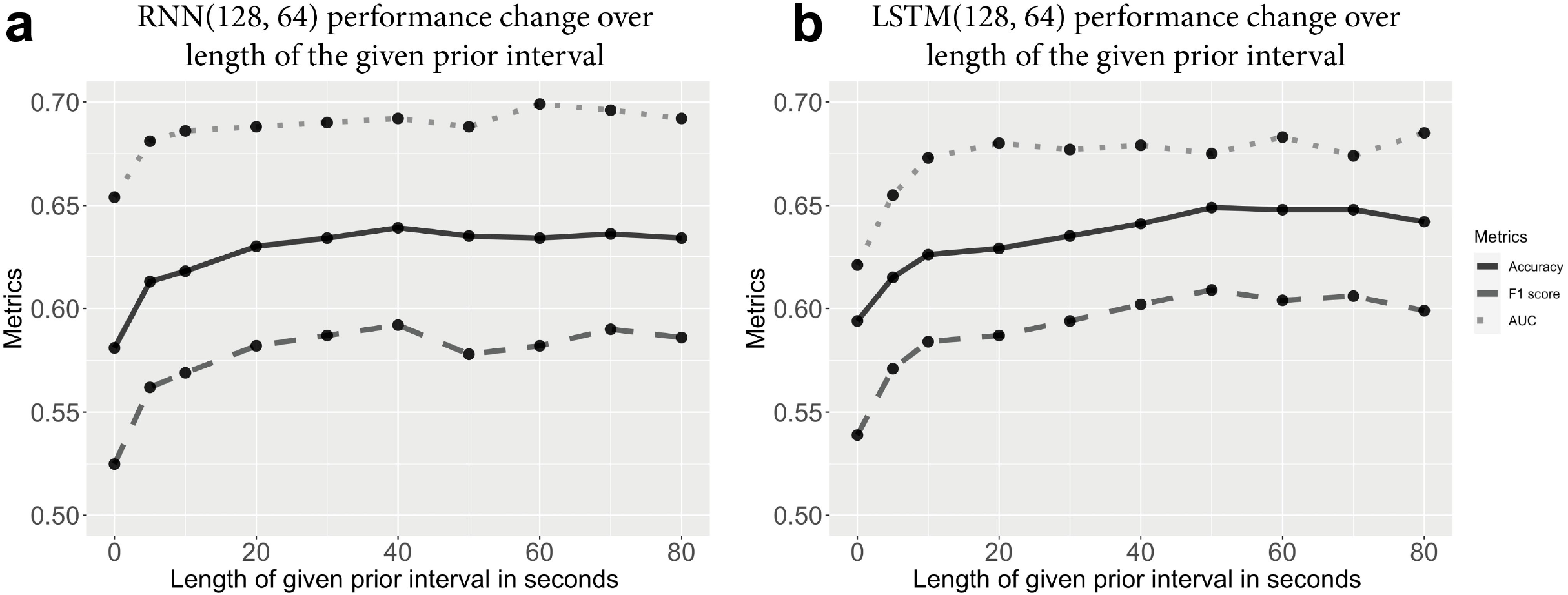
Model performance change over the length of a given prior interval. Model performance is evaluated using validation accuracy, AUC and F1 score. The length of prior intervals used are 0 (0s), 5 (150s), 10 (300s), 20 (600s), 30 (900s), 40 (1200s), 50 (1500s), 60 (1800s), 70 (2100s) and 80 (2400s).

To summarize, the inclusion of history accelerometry measures through sequential models has improved the model performance compared to non-sequential models. Overall, both RNN and LSTM achieved their best prediction accuracy at b around 50, indicating that the prior 250 seconds of movements are highly informative for the current sleep stage.

### 3. The Contribution of Variance and Range as Features

In addition to features that were directly available from the preprocessing output, we found that the presence of two additional derived variables has strongly improved the model performance in all means (**Supplement Table A.2**). The two features are the local range and variance of each activity metric (angle x, y, z, and ENMO) from the left or right wrist within each 30-second interval. Intuitively, the range and variance reflect the first and second moments of the distribution of the time-varying accelerometry measurements, which could be informative for the different movement patterns when participants were at various sleep stages. The ‘heuristic algorithm’ in Van Hees et al. (2019) determined sleep window heavily based upon the local variance of angle-z accelerations. However, we believed that simply thresholding by angle-z accelerations might be non-specific and insufficient for the task of sleep stage classification due to device rotation and change of position. Hence, we experimented with expanding our feature set to include the variance and range of all the three-axis accelerations and the overall intensities. Under the optimally trained model structure of LSTM 128&64, as shown in **Figure 3**, adding either one of them into the model greatly improved the performance in all the evaluation metrics. In particular, the average validation accuracy increased from 58% to 65%, the average F1 score increased from 0.51 to 0.58, and AUC increased from 0.60 to 0.67. Furthermore, models with the two additional features also relaxed the requirement of b and allowed the inclusion of a shorter historical window to achieve the same optimal results.

**Figure 3.**
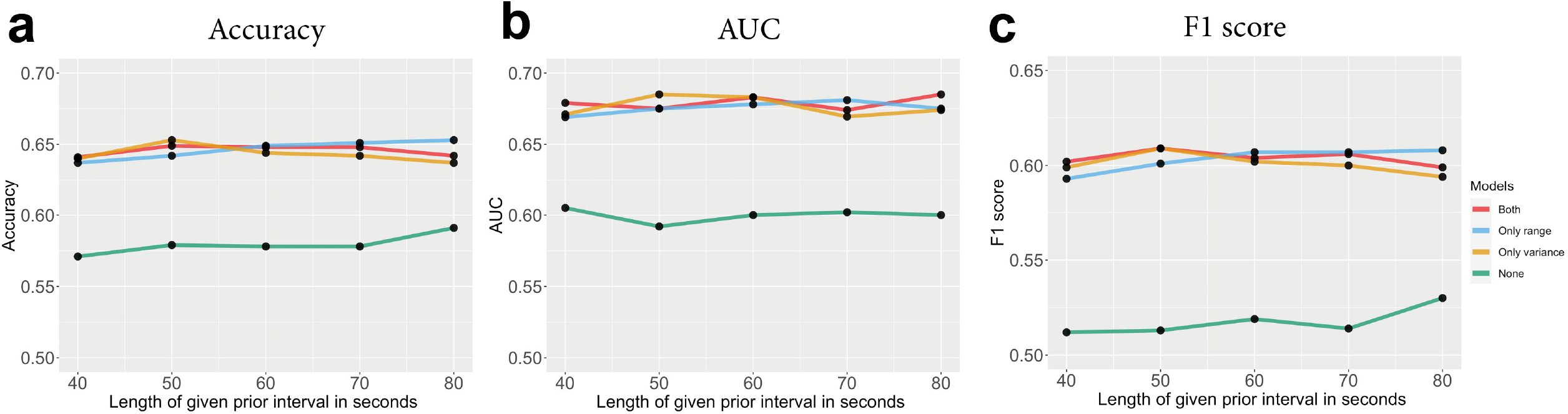
Model performance with/without variance and range of all the three-axis measures. LSTM with 2 hidden layers of 128 and 64 nodes is used as the testing model. Model performance is evaluated using validation accuracy, AUC and F1 score.

### 4. The Impact of Clinical Characteristics on Prediction Performance

As noted in the Method section, our algorithm was trained upon data collected from sleep clinic visitors and 75% (21 out of 28) were diagnosed with at least one of the following sleep disorders: idiopathic hypersomnia, Restless legs, sleep apnoea, narcolepsy, sleep paralysis, obstructive sleep apnoea, RBD insomnia and insomnia. In addition to assessing the average model performance across the entire sample, we are also interested in evaluating the applicability of the trained model on individual participants and understand the impact of underlying sleep conditions.

To evaluate the subject heterogeneity, we calculated the individual’s cross-validated model performance under the established optimal model structure (LSTM 128&64, b=50, all features) for each participant and compared them by participants’ clinical characteristics (**Supplement Table A.3**). As shown in **A.3**, even though the training accuracies achieved around 68% for all participants, the validation accuracies reveal a wide range (35%-81% with a median of 67.8%). For 9 out 27 participants (33%), our model achieved more than 70% validation accuracies, yet the model failed to be generalized to another 4 participants who are all diagnosed with certain sleep disorders (validation accuracies less than 50%). **Figure 4** demonstrates the distribution of the three evaluation metrics (accuracy, F1, AUC) over age, gender and disease status. Participants with sleep disorders tend to be older in age compared to normal participants. Although the overall model performances are comparable between sleep disorder patients and normal participants, we observed a higher variability in validation accuracies among diseased participants as compared to normal participants (p=0.08 using Bartlett’s test for equal variance, standard deviation and interquartile range in **Table 2**). In addition, the four participants with the lowest prediction accuracies are those with serious disorders (i.e., restless leg, restless nocturia, obstructive sleep apnoea and mild sleep apnoea), indicating that the population model might not be sufficient in characterizing abnormal sleep patterns and might require larger training samples of similar pathologies.

**Figure 4.**
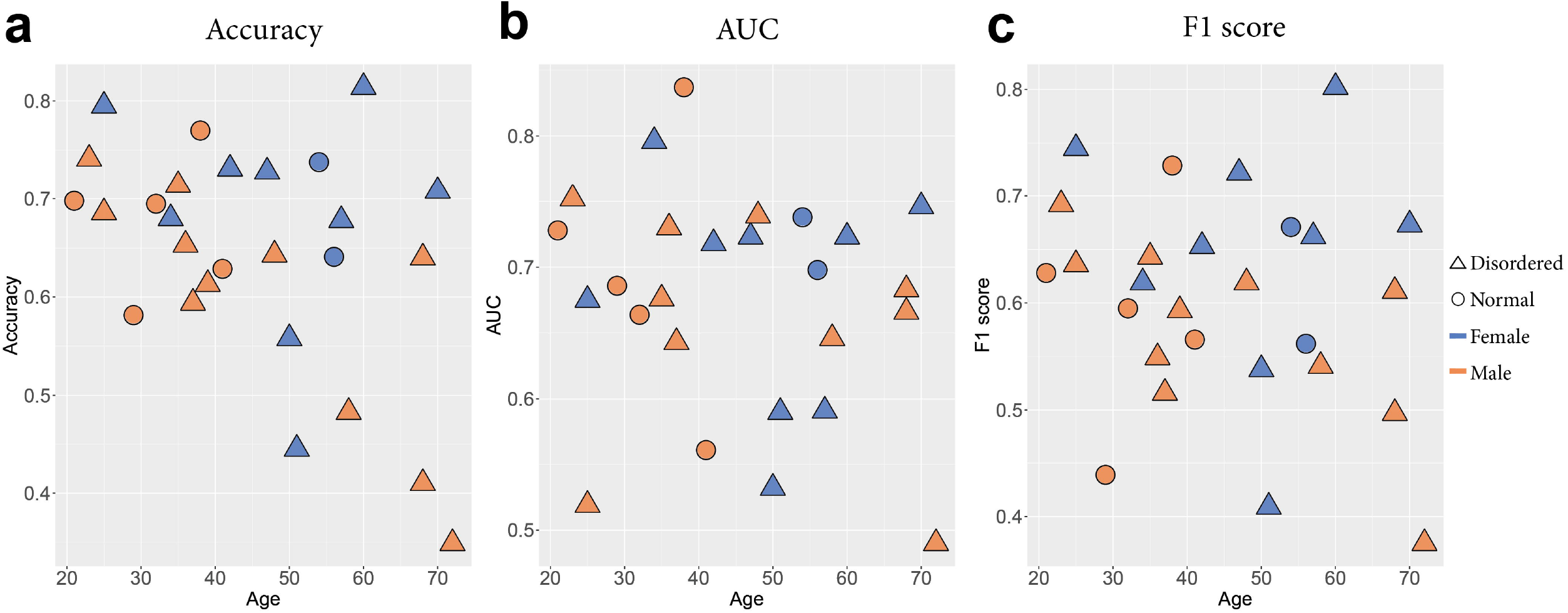
Distribution of accuracy, AUC and F1 score over three factors: age, gender and disorder status.

**Table 2:** Stand Deviation (SD) and Interquartile Range (IQR) of Accuracy, F1 score and AUC in normal and diseased population.

| Metrics | Validation accuracy |  | F1 score |  | AUC |  |
| --- | --- | --- | --- | --- | --- | --- |
|  | SD | IQR | SD | IQR | SD | IQR |
| Normal | 0.065 | 0.083 | 0.092 | 0.085 | 0.083 | 0.058 |
| Diseased | 0.130 | 0.145 | 0.109 | 0.128 | 0.086 | 0.110 |

## MATERIALS AND METHODS

In this section, we introduce the PSG sleep data, the input features, the algorithms and model evaluation metrics used in this study.

### 1. Dataset

We used data from the publicly available Newcastle PSG+Accelerometer study as available for download^21^. The dataset contained one-night polysomnography (PSG) assessment from 28 adult participants at the Freeman Hospital in the UK. These data were previously processed and used as the gold standard to validate existing sleep detection algorithms ^22^. During the monitor period of the study, bilateral eye movements, leg movements, respiratory and sleep electroencephalogram (EEG) were recorded for each participant. Their PSG signals and respiratory events were then used to score sleep stages into one of the five categories including awakening, rapid eye movement (REM) sleep and N1, N2 and N3 non-REM in 30-second resolution. Among the 28 patients, 8 of them were healthy normal and the remaining 21 were diagnosed with at least one of the following sleep disorders: idiopathic hypersomnia, restless legs, narcolepsy, REM sleep behavior disorder (RBD), insomnia and sleep apnoea. Data from one participant with a sleep disorder was discarded due to the absence of records from one wrist.

Meanwhile, the participants wore two accelerometers from brand GENEActiv on both left and right wrists. The devices recorded time series of 3-axis accelerations (x, y, z coordinates relative to the accelerometers) during the wear time at a frequency of 85.7Hz as a measure of the continuous levels of physical activity during the participants’ wrist movements. Additionally, the ambient temperature was also tracked overtime to assist non-wear detection. The raw features were stored in separate binary files and processed using the standard pipeline for GENEActiv data available at https://github.com/wadpac/psg-ncl-acc-spt-detection-eval and R package ‘GGIR’^22,23^ and converted into csv files. In particular, the preprocessing steps auto-calibrated local gravity and aggregated the densely measured raw signals into time series of 5-second epoch length (one value per 5s per feature). We extracted two sets of main features from the raw data: the corresponding angles (in degrees) of each of the axes relative to the horizontal plane; and Euclidean Norm Minus One (ENMO) ^24^ as a one-dimensional summary that characterizes the concurrent magnitude of physical activity intensities. Unlike the raw 3-axis accelerations, ENMO is invariant to the device orientation and has been reported to be particularly sensitive to movements with low-intensity levels. It is calculated by:

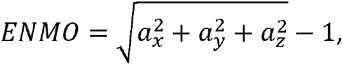

where *a_x_*, *a_y_*, *a_z_* are the median values of raw acceleration in x, y and z coordinates within each 5 second epoch. Other covariates include participants’ gender, age and sleep disorder diagnosis. We summarized the data type and distribution of the participants’ clinical characteristics, the raw features collected using accelerometers and outcome variables in **Table 3**.

**Table 3:** Description of preprocessed raw variables from the original dataset.

| Feature Name | Unit | Time dependent | Mean<br>(Standard deviation) |
| --- | --- | --- | --- |
| Age | years | No | 45.011 (14.397) |
| Gender | female / male | No | 11 females, 16 males |
| Disorder | normal / disorder | No | 7 normal, 20 disordered |
| Left arm temperature | celsius degree (°C) | Yes | 31.005 (2.303) |
| Acceleration in left arm angle x | g-units (1g = 1000mg) | Yes | -2.148 (40.011) |
| Acceleration in left arm angle y | g-units (1g = 1000mg) | Yes | 10.139 (29.743) |
| Acceleration in left arm angle z | g-units (1g = 1000mg) | Yes | -10.775 (42.324) |
| Right arm temperature | celsius degree (°C) | Yes | 32.12 (2.582) |
| Acceleration in right arm angle x | g-units (1g = 1000mg) | Yes | -6.453 (40.435) |
| Acceleration in right arm angle y | g-units (1g = 1000mg) | Yes | -1.401 (30.557) |
| Acceleration in right arm angle z | g-units (1g = 1000mg) | Yes | -15.667 (41.271) |
| Sleep stage (outcome) | wake / REM / non-REM | No | Wake:55464(30.5%)<br>REM: 21306(11.7%)<br>Non-REM:104850(57.8%) |

In addition to the direct outputs from the preprocessing pipelines, we also generated new features and tested whether adding them to the model would improve the performance. As demonstrated in **Figure 1a**, since the measurements from wrist-worn accelerometers were assessed every 5 seconds while the true sleep stage was detected every 30 seconds, we also calculated the log-transformed variances and ranges of ENMO, *a_x_*, *a_y_* and *a_z_* within each 30-s window to reflect characteristics of the higher-order moments of the original features. All the continuous features are normalized before model fitting by subtracting the mean then divided by the standard deviation.

### 2. Model

We mainly focused on developing sequential deep learning models such as RNN and LSTM Network to classify sleep versus awake, and further differentiate REM and non-REM within the sleep windows. RNN, which was originally designed for nature language analysis, is a suitable model for time series. Many studies show that RNN often achieves superior performance in dealing with time-series data compared to other models. In order to incorporate temporal dependence from both the input and output variables, a parameter called “window size” was used to control the amount of historical data to include for predicting the current sleep stage. By doing so, RNN-based methods allow the model to learn the information in sequential order, and automatically detect the additional pattern behind the sequential data that might otherwise be neglected if the accelerometry features were treated as independent. However, when the window size of the past records becomes larger, conventional RNN becomes incapable of handling such long-term dependencies due to gradient vanishing and gradient explosion^25^. Hence, we also deployed LSTM as a special variant of RNN that is able to accommodate long-term dependencies. In particular, as shown in **Figure 1b**, the outcome data are sleep stage labels in 30s resolution, while the input features were collected every 5s epoch length. We developed our algorithm in python using ‘torch’ package and implemented in AWS cloud computing service with one core. The final package is available on the github repository.

We compared RNN and LSTM models with several existing classifiers such as logistic regression, Perceptron, Extra Tree, and Random Forest that have previously been reported^19^ to achieve good performance in similar tasks of separating the awake and asleep states. However, neither of the aforementioned ML models was originally designed for time series data. Hence when applying to our data, we treated the sequential data within each 30-second window as independent features.

### 3. Evaluation Metrics

For each method, we used the leave-one-subject-out cross-validation to train and evaluate the model. We assessed and compared the model performances for multi-class classification using the conventional metrics, including average training and validation accuracy, weighted precision, recall, F1 score in the test data. Area-under-curve (AUC) was also calculated for logistic regression, RNN and LSTM that produce continuous probability estimation. In particular, prediction accuracy was calculated as the proportion of the correct predictions across all classes among the total number of predictions. For metrics that are originally defined for the binary classification task, i.e., precision, recall, F1 score, and AUC, we treated the data as a collection of binary problems, one for each class, then calculated these metrics for each class and reported the average weighted by the number of true instances in each class. For example, the multi-class precision is calculated as:

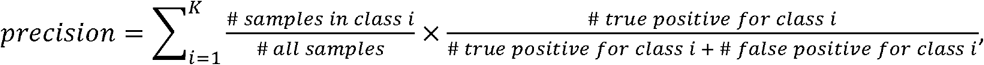

where is K is the number of classes in the data. The weighted metrics incorporated the proportions of true instances in each label and accounted for label imbalances in the training and testing data. All of these metrics were calculated using ‘sklearn.metrics’ in Python^26^.

## DISCUSSION

Actigraphy data have been long used to record movement in sleep and wake. However, autonomous algorithms in differentiating sleep and wake often require large training samples. Furthermore, fine classification of detailed sleep stages still relies heavily upon experts’ inspection of multiple signals from movement, respiratory rate and heart rate variability collected under professional settings. Given the increasing availability of wearable computing sensors that allow in-home signal tracking, it is of great interest to examine the extent to which actigraphy data alone could detect and classify sleep using advanced machine learning algorithms. The challenges in such tasks come from both the high throughput input variables as well as the temporal dependency in the recorded measurements. Not being able to account for neighboring information often leads to fragmented classification labels with low accuracy.

We have compared a comprehensive list of conventional (e.g., logistic regression, support vector machine and random forest) and modern neural network-based (RNN and LSTM) machine learning methods for the task of classifying wake, REM and non-REM sleep using a relatively small dataset. The input features for our prediction are only limited to data from the wrist-worn devices including three-axis accelerations, angles and temperature. Among all the algorithms, the LSTM-based models that enable the information borrowed from previous records and take advantage of the underlying deep dependency structures achieved the best performance. We have identified the optimal tuning parameters and model structures using leave-one-subject out cross-validation and demonstrated that feeding 50 prior intervals of history records resulted in the best prediction accuracy. We have also shown that, in addition to the raw acceleration features, adding derived local feature variability (range and variance) could greatly improve model performance in almost all the performance metrics. For example, we have observed about 12% increase in average validation accuracy, 13.7% increase in average F1 score and 11% increase in AUC. We have demonstrated that actigraphy data alone, when combined with adequate feature sets and powerful LSTM modeling techniques, are useful in differentiating wake, REM and non-REM sleep by achieving 65% average accuracies compared to the reference accuracy of 57%. As expected, our average performance using only accelerometry data is inferior to models trained with complex multimodal biosignals and on large normative samples. However, we believe part of the reasons are due to limited sample size and lack of normal subjects in our training data. In fact, our algorithm was able to achieve close to optimal performance on a subset of the individuals with the highest validation accuracy achieving 81% and 33% participants having higher than 70% accuracies. Such results might provide practical insights on the utility of accelerometry data for capturing important sleep characteristics with relatively lower cost. It might also allow for the extraction of sleep-related features in large cohort studies with fast and reproducible methods. The final software for the developed model is available on the github repository at: https://github.com/jianhuupenn/Sleep-stage-classification.

We would like to emphasize that our paper is different from existing literature on sleep classification in that the goal is not to achieve the best prediction accuracy. Instead, we wish to examine the essential elements for a useful classification model for accelerometry data through a set of experiments. We have concluded that although it is possible to classify sleep stages with reasonable performance using only accelerometry variables and a small sample, it is crucial to utilize a model that accounts for temporal dependency and incorporates the local variability of the features. Including sufficient training data of subjects with sleep disorders is also important for the model to be generalized to external data. There remain several directions of improvements for the current algorithm. One of them, as indicated by the post-hoc analysis that correlated model performances with subjects’ characteristics, is that the model performances were on average worse among those with sleep disorders compared to normal subjects.

They also tend to be and more variable due to the heterogeneous diagnosis in the training sample. Similar phenomenon might also be true for existing sleep classification algorithms as the diagnostic of sleep disorders are not often available as part of the training data. Future algorithms could be improved by including samples of the diseased populations with homogeneous diagnoses and respecting differential sleep characteristics among normal subjects and sleep disorder patients. Additionally, one could further incorporate neighborhood correlation from both historical data and future data such as consider implementing LSTM models with bi-directional layers of forward and backward dependencies. We did not implement such models due to constraints on the size of training data^27^.

## Data and code availability

The published data sets from Newcastle PSG study used in this manuscript are available through the following websites: http://doi.org/10.5281/zenodo.1160410

Code for this analysis is available at: https://github.com/jianhuupenn/Sleep-stage-classification

