## Supplementary Materials for "Three-level Sleep Stage Classification Based on Wrist-worn Accelerometry Data Alone"

**Supplementary Tables for ‘Three-level Sleep Stage Classification based on Wrist-worn Accelerometry Data Alone’**

**Table A.1** Prediction performances for 3-level sleep stage classification using conventional machine learning models and under various lengths of past data and model complexity of RNN

| Model | Training accuracy | Validation accuracy | Precision | Recall | F1 score | AUC |
| --- | --- | --- | --- | --- | --- | --- |
| LR | **0.648** | **0.631** | **0.572** | **0.631** | **0.571** | **0.656** |
| Perceptron | 0.544 | 0.535 | 0.577 | 0.535 | 0.477 | NA |
| Extra tree | 1.0 | 0.455 | 0.531 | 0.455 | 0.466 | NA |
| RF | 1.0 | 0.594 | 0.577 | 0.594 | 0.565 | NA |
| --- | --- | --- | --- | --- | --- | --- |
| RNN(b=0, 64_32) | 0.625 | 0.576 | 0.580 | 0.576 | 0.525 | 0.648 |
| RNN(b=5, 64_32) | 0.631 | 0.585 | 0.592 | 0.545 | 0.535 | 0.654 |
| RNN(b=10, 64_32) | 0.631 | **0.604** | 0.578 | 0.604 | **0.550** | 0.651 |
| RNN(b=20, 64_32) | 0.630 | 0.586 | 0.588 | 0.586 | 0.535 | 0.655 |
| RNN(b=30, 64_32) | 0.631 | 0.591 | 0.590 | 0.593 | 0.540 | 0.656 |
| RNN(b=40, 64_32) | 0.633 | 0.599 | 0.593 | 0.599 | 0.549 | **0.658** |
| RNN(b=0, 128_64) | 0.628 | 0.581 | 0.579 | 0.581 | 0.525 | 0.654 |
| RNN(b=5, 128_64) | 0.648 | 0.613 | 0.604 | 0.613 | 0.562 | 0.681 |
| RNN(b=10, 128_64) | 0.654 | 0.618 | 0.616 | 0.618 | 0.569 | 0.686 |
| RNN(b=20, 128_64) | 0.657 | 0.630 | 0.620 | 0.630 | 0.582 | 0.688 |
| RNN(b=30, 128_64) | 0.657 | 0.634 | 0.621 | 0.635 | 0.587 | 0.690 |
| RNN(b=40, 128_64) | **0.659** | **0.639** | **0.624** | **0.639** | **0.592** | **0.692** |
| RNN(b=50, 128_64) | 0.662 | 0.635 | 0.628 | 0.635 | 0.578 | 0.688 |
| RNN(b=60, 128_64) | 0.666 | 0.634 | 0.628 | 0.634 | 0.582 | 0.699 |
| RNN(b=70, 128_64) | 0.664 | 0.636 | 0.646 | 0.636 | 0.590 | 0.696 |
| RNN(b=80, 128_64) | 0.667 | 0.634 | 0.634 | 0.634 | 0.586 | 0.692 |
| RNN(b=50, 256_128) | 0.670 | 0.603 | 0.637 | 0.603 | 0.555 | 0.691 |
| RNN(b=60, 256_128) | 0.671 | 0.623 | 0.629 | 0.623 | 0.575 | 0.701 |
| RNN(b=70, 256_128) | 0.660 | 0.631 | 0.626 | 0.631 | 0.580 | 0.686 |
| RNN(b=80, 256_128) | 0.661 | 0.624 | 0.623 | 0.624 | 0.573 | 0.683 |

**Table A.2** Impact on model performance by including variance and range as features in LSTM under various model complexity.

| Model | Feature | Backward | Forward | Training accuracy | Validation accuracy | Precision | Recall | F1 score | AUC |
| --- | --- | --- | --- | --- | --- | --- | --- | --- | --- |
| LSTM_128_64 | Both | 0 | 0 | 0.615 | 0.594 | 0.567 | 0.594 | 0.539 | 0.621 |
| LSTM_128_64 | Both | 5 | 0 | 0.645 | 0.615 | 0.65 | 0.614 | 0.571 | 0.655 |
| LSTM_128_64 | Both | 10 | 0 | 0.654 | 0.626 | 0.616 | 0.626 | 0.584 | 0.673 |
| LSTM_128_64 | Both | 20 | 0 | 0.663 | 0.629 | 0.611 | 0.629 | 0.587 | 0.680 |
| LSTM_128_64 | Both | 30 | 0 | 0.671 | 0.635 | 0.618 | 0.635 | 0.594 | 0.677 |
| LSTM_128_64 | Both | 40 | 0 | 0.677 | 0.641 | 0.622 | 0.641 | 0.602 | 0.679 |
| LSTM_128_64 | Both | 50 | 0 | 0.681 | **0.649** | 0.621 | 0.649 | **0.609** | 0.675 |
| LSTM_128_64 | Both | 60 | 0 | 0.683 | 0.648 | 0.626 | 0.648 | 0.604 | 0.683 |
| LSTM_128_64 | Both | 70 | 0 | 0.684 | 0.648 | 0.630 | 0.648 | 0.606 | 0.674 |
| LSTM_128_64 | Both | 80 | 0 | 0.685 | 0.642 | 0.625 | 0.642 | 0.599 | **0.685** |
| --- |  |  |  |  |  |  |  |  |  |
| LSTM_256_128 | Both | 0 | 0 | 0.619 | 0.595 | 0.570 | 0.595 | 0.541 | 0.636 |
| LSTM_256_128 | Both | 5 | 0 | 0.654 | 0.612 | 0.607 | 0.612 | 0.569 | 0.677 |
| LSTM_256_128 | Both | 10 | 0 | 0.665 | 0.611 | 0.623 | 0.611 | 0.571 | 0.687 |
| LSTM_256_128 | Both | 20 | 0 | 0.675 | 0.627 | 0.634 | 0.627 | 0.591 | 0.687 |
| LSTM_256_128 | Both | 30 | 0 | 0.681 | 0.630 | 0.626 | 0.630 | 0.592 | 0.684 |
| --- |  |  |  |  |  |  |  |  |  |
| LSTM_128_64 | None | 40 | 0 | 0.618 | 0.571 | 0.552 | 0.571 | 0.512 | 0.605 |
| LSTM_128_64 | None | 50 | 0 | 0.619 | 0.579 | 0.555 | 0.579 | 0.513 | 0.592 |
| LSTM_128_64 | None | 60 | 0 | 0.624 | 0.578 | 0.564 | 0.578 | 0.519 | 0.600 |
| LSTM_128_64 | None | 70 | 0 | 0.626 | 0.578 | 0.565 | 0.578 | 0.514 | 0.602 |
| LSTM_128_64 | None | 80 | 0 | 0.629 | 0.591 | 0.571 | 0.591 | 0.530 | 0.600 |
| --- |  |  |  |  |  |  |  |  |  |
| LSTM_128_64 | Only range | 40 | 0 | 0.673 | 0.637 | 0.614 | 0.637 | 0.593 | 0.669 |
| LSTM_128_64 | Only range | 50 | 0 | 0.676 | 0.642 | 0.618 | 0.642 | 0.601 | 0.675 |
| LSTM_128_64 | Only range | 60 | 0 | 0.678 | 0.649 | 0.620 | 0.649 | 0.607 | 0.678 |
| LSTM_128_64 | Only range | 70 | 0 | 0.680 | 0.651 | 0.624 | 0.651 | 0.607 | 0.681 |
| LSTM_128_64 | Only range | 80 | 0 | 0.680 | 0.653 | 0.618 | 0.653 | 0.608 | 0.675 |
| --- |  |  |  |  |  |  |  |  |  |
| LSTM_128_64 | Only var | 40 | 0 | 0.673 | 0.640 | 0.616 | 0.640 | 0.599 | 0.671 |
| LSTM_128_64 | Only var | 50 | 0 | 0.676 | 0.653 | 0.632 | 0.653 | 0.609 | 0.685 |
| LSTM_128_64 | Only var | 60 | 0 | 0.677 | 0.644 | 0.617 | 0.644 | 0.602 | 0.683 |
| LSTM_128_64 | Only var | 70 | 0 | 0.679 | 0.642 | 0.622 | 0.642 | 0.600 | 0.669 |
| LSTM_128_64 | Only var | 80 | 0 | 0.679 | 0.637 | 0.6133 | 0.637 | 0.594 | 0.674 |
| --- |  |  |  |  |  |  |  |  |  |
| LSTM_128_64 | Both | 40 | 0 | 0.677 | 0.641 | 0.622 | 0.641 | 0.602 | 0.679 |
| LSTM_128_64 | Both | 40 | 5 | 0.676 | 0.643 | 0.640 | 0.643 | 0.600 | 0.668 |
| LSTM_128_64 | Both | 40 | 10 | 0.674 | 0.641 | 0.625 | 0.641 | 0.598 | 0.654 |
| LSTM_128_64 | Both | 40 | 20 | 0.673 | 0.640 | 0.616 | 0.640 | 0.594 | 0.650 |
| LSTM_128_64 | Both | 40 | 30 | 0.672 | 0.642 | 0.616 | 0.642 | 0.594 | 0.631 |
| LSTM_128_64 | Both | 40 | 40 | 0.671 | 0.644 | 0.620 | 0.644 | 0.598 | 0.626 |

**Table A.3** Detailed cross-validated performance metrics by individuals

| ID | Disorder | Training accuracy | Validation accuracy | Precision | Recall | F1 score | AUC |
| --- | --- | --- | --- | --- | --- | --- | --- |
| 1 | 1 | 0.674 | 0.795 | 0.756 | 0.795 | 0.745 | 0.675 |
| 2 | 0 | 0.679 | 0.698 | 0.610 | 0.698 | 0.628 | 0.728 |
| 14 | 1 | 0.692 | 0.349 | 0.543 | 0.349 | 0.375 | 0.490 |
| 17 | 1 | 0.679 | 0.728 | 0.722 | 0.728 | 0.722 | 0.723 |
| 21 | 1 | 0.673 | 0.814 | 0.809 | 0.814 | 0.802 | 0.723 |
| 23 | 1 | 0.674 | 0.708 | 0.665 | 0.708 | 0.673 | 0.746 |
| 27 | 0 | 0.675 | 0.770 | 0.776 | 0.770 | 0.729 | 0.837 |
| 28 | 1 | 0.688 | 0.483 | 0.872 | 0.483 | 0.541 | 0.646 |
| 29 | 1 | 0.691 | 0.445 | 0.475 | 0.445 | 0.409 | 0.590 |
| 31 | 1 | 0.683 | 0.686 | 0.596 | 0.686 | 0.636 | 0.519 |
| 32 | 1 | 0.677 | 0.640 | 0.586 | 0.640 | 0.611 | 0.666 |
| 34 | 1 | 0.682 | 0.678 | 0.648 | 0.678 | 0.662 | 0.591 |
| 35 | 1 | 0.688 | 0.613 | 0.695 | 0.613 | 0.593 | NA |
| 38 | 0 | 0.682 | 0.629 | 0.516 | 0.629 | 0.566 | 0.561 |
| 39 | 1 | 0.674 | 0.731 | 0.598 | 0.731 | 0.653 | 0.718 |
| 42 | 1 | 0.687 | 0.653 | 0.491 | 0.653 | 0.549 | 0.730 |
| 45 | 1 | 0.681 | 0.558 | 0.525 | 0.558 | 0.538 | 0.532 |
| 48 | 1 | 0.690 | 0.410 | 0.757 | 0.410 | 0.497 | 0.683 |
| 49 | 1 | 0.682 | 0.643 | 0.777 | 0.643 | 0.619 | 0.739 |
| 50 | 0 | 0.686 | 0.641 | 0.502 | 0.641 | 0.562 | 0.699 |
| 51 | 1 | 0.684 | 0.680 | 0.580 | 0.680 | 0.619 | 0.796 |
| 52 | 1 | 0.680 | 0.741 | 0.760 | 0.741 | 0.692 | 0.752 |
| 53 | 1 | 0.678 | 0.714 | 0.600 | 0.714 | 0.643 | 0.676 |
| 56 | 1 | 0.679 | 0.594 | 0.639 | 0.594 | 0.516 | 0.643 |
| 57 | 0 | 0.683 | 0.738 | 0.620 | 0.738 | 0.671 | 0.738 |
| 59 | 0 | 0.687 | 0.582 | 0.357 | 0.582 | 0.439 | 0.686 |
| 60 | 0 | 0.674 | 0.695 | 0.530 | 0.695 | 0.595 | 0.664 |
